# Combining serological and contact data to derive target immunity levels for achieving and maintaining measles elimination

**DOI:** 10.1101/201574

**Authors:** Sebastian Funk, Jennifer K. Knapp, Emmaculate Lebo, Susan E. Reef, Alya J. Dabbagh, Katrina Kretsinger, Mark Jit, W. John Edmunds, Peter M. Strebel

## Abstract

**Background:** Vaccination has reduced the global incidence of measles to the lowest rates in history. However, local interruption of measles virus transmission requires sustained high levels of population immunity that can be challenging to achieve and maintain. The herd immunity threshold for measles is typically stipulated at 90–95%. This figure does not easily translate into age-specific immunity levels required to interrupt transmission. Previous estimates of such levels were based on speculative contact patterns based on historical data from high-income countries. The aim of this study was to determine age-specific immunity levels that would ensure elimination of measles when taking into account empirically observed contact patterns.

**Methods:** We combined estimated immunity levels from serological data in 17 countries with studies of age-specific mixing patterns to derive contact-adjusted immunity levels. We then compared these to case data from the 10 years following the seroprevalence studies to establish a contact-adjusted immunity threshold for elimination. We lastly combined a range of hypothetical immunity profiles with contact data from a wide range of socioeconomic and demographic settings to determine whether they would be sufficient for elimination.

**Results:** We found that contact-adjusted immunity levels were able to predict whether countries would experience outbreaks in the decade following the serological studies in about 70% of countries. The corresponding threshold level of contact-adjusted immunity was found to be 93%, corresponding to an average basic reproduction number of approximately 14. Testing different scenarios of immunity with this threshold level using contact studies from around the world, we found that 95% immunity would have to be achieved by the age of five and maintained across older age groups to guarantee elimination. This reflects a greater level of immunity required in 5–9 year olds than established previously.

**Conclusions:** The immunity levels we found necessary for measles elimination are higher than previous guidance. The importance of achieving high immunity levels in 5–9 year olds presents both a challenge and an opportunity. While such high levels can be difficult to achieve, school entry provides an opportunity to ensure sufficient vaccination coverage. Combined with observations of contact patterns, further national and sub-national serological studies could serve to highlight key gaps in immunity that need to be filled in order to achieve national and regional measles elimination.

## Background

Measles, a highly contagious immunising infection, could be a future target for eradication. [1, 2] Since the introduction of vaccination in the early 1960s, mortality and morbidity from measles has declined drastically. [3] Nevertheless, outbreaks continue to occur, and achieving regional elimination, or interruption of transmission, has been challenging. [4]

Control of measles is achieved through vaccination in early childhood, and the vaccine is part of routine immunisation schedules worldwide. In principle, a functioning health system would aim to vaccinate every child. In practice 100% coverage with all recommended doses is never achieved. Moreover, not every administration of a vaccine confers immunity, and protection from a vaccine can wane over time. However, even if not everyone in a population is immune, the indirect protection provided by the presence of immune individuals can be sufficient to prevent outbreaks. [5] For measles, it has been shown that in a randomly mixing population the level of immunity required to achieve this so-called “herd immunity” is in the order of 90-95%. [6]

Knowledge of the level of immunity required in a population to achieve herd immunity can be used to set national vaccination targets. However, even if current levels of vaccination are high enough to achieve the level of immunisation required for herd immunity, outbreaks can occur if there are immunity gaps in older age groups. To assess the ability of a country or region to achieve and maintain elimination, that is the sustained absence of endemic transmission, immunity levels must therefore be considered across all age groups. These levels are affected by historical and current routine vaccination coverage, but also by vaccination campaigns and past outbreaks that conferred natural immunity.

For this reason, in the late 1990s, the World Health Organization (WHO) European Region (EURO) derived age-specific target immunity profiles, or the levels of immunity necessary in different age groups in order to achieve elimination. [7] These profiles are widely applied within and occasionally outside Europe to assess progress towards elimination. [8, 9, 10, 11, 12, 13, 14, 15, 16] Based on a basic reproduction number (or number of secondary cases produced by a typical infective in a totally susceptible population) of 11, it was recommended to ensure that at least 85% of 1–4 year olds, 90% of 5–9 year olds and 95% of 10 year olds and older possess immunity against measles. [17] Unlike vaccination coverage targets, immunity targets reflect the effect of susceptibility in all age groups and highlight the potential need for campaigns to close any gaps in immunity.

The aforementioned target immunity levels derived in the late 1990s were based on assumed age-specific contact patterns based on pre-vaccination measles epidemiology in England and Wales. Since then, much work has gone into better quantifying the amount of transmission-relevant contact occurring between different age groups. Diary-based studies have been conducted across Europe [18, 19], as well as in Vietnam [20] China [21], Uganda [22], Zimbabwe [23] and elsewhere. While other methods for measuring social contact patterns exist [24, 25, 26], contact data from diary studies have become the de facto standard for studying age-specific infectious disease dynamics. Mathematical models of transmission based on these observed patterns have consistently outperformed those based on homogeneous mixing. [27, 28, 29]

Here, we aimed to evaluate current guidelines on target immunity levels for measles taking into account contact patterns observed in diary studies. To this end, we combined the observed age-specific social mixing patterns with observed or hypothesised immunity levels to calculate a *contact-adjusted* immunity, akin to the mean level of immunity across the population but taking into account that some age groups have more contact with each other than others. We validated this method by testing the extent to which contact-adjusted immunity levels based on nationwide serological studies conducted in different countries in the late 1990s and early 2000s could have been used to predict the case load in the following decade. We then calculated contact-adjusted immunity levels from a range of hypothetical scenarios of age-specific immunity, including previous recommended immunity levels. We assessed whether these levels would be sufficient for achieving and maintaining elimination.

## Methods

### Predicting elimination from seroprevalence data

We estimated population-level immunity levels from seroprevalence data using the different model variants outlined below and compared them to the number of cases experienced over 10 years using Spearman’s rank correlation coefficient.

We further tested different thresholds for these levels to classify countries as being at risk of outbreaks or not. We calculated the misclassification error (MCE) as the proportion of countries that were incorrectly classified based on the given immunity threshold level and a threshold of the number of cases experienced in the 10 years following the seroprevalence study.

### Immunity model: contact-adjusted vs. plain

We assume that the force of infection *λ_i_* experienced by age group *i* only depends on the rate of contact with the same and other age groups and the prevalence of infection in the respective age groups:

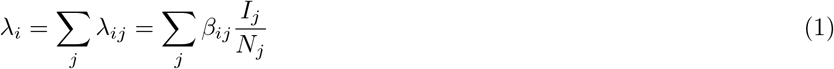

where *λ_ij_* is the force of infection exerted by age group *j* on age group *i*, *β_ij_* is the infection rate, or the rate at which individuals in age group *i* contact individuals out of a total number *N*_*j*_ in age group *j* and become infected if these are infectious, and *I*_*j*_ is the number of infectious people in age group *j*. This formulation of the force of infection assumes that the rate of infection between two random individuals depends on their ages only, and that the probability of a given contacted member of age group *j* to be with someone infectious depends on population-level prevalence of infection only.

We further write the infection rate *β_ij_* as

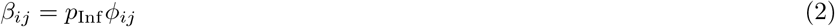

where *p*_Inf_ is the probability that a contact between a susceptible and infectious person leads to infection, here assumed age-independent, and *φ_ij_* is the number of contacts an individual of age group *j* makes with those of age group *i* per unit time.

The basic reproduction number *R*_0_ is defined as the mean number of new cases generated by a single infectious individual in a completely susceptible population. In a system with multiple host types (here: age groups), it can be calculated as the spectral radius (or largest eigenvalue) of the next-generation matrix (NGM) **K** [30]

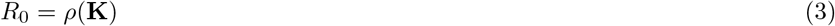

The elements of the next-generation matrix **K** can be written as

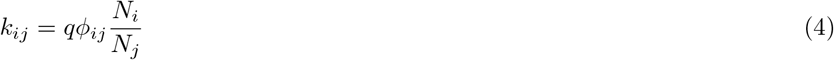

where *q* is a scale factor that, assuming that infectiousness stays constant while a person is infectious, is the probability of infection upon contact *p*_Inf_ multiplied with the duration of infectiousness *D*_Inf_. If a proportion *r*_*i*_ of age group *i* is immune, this changes the initially susceptible population from *N*_*i*_ to *N*_*i*_(1 *−r_i_*). The reproduction number for an invading infection in such a population is

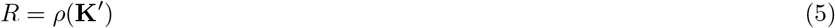

where, again, *ρ* denotes the spectral radius and **K**′ is a matrix with elements

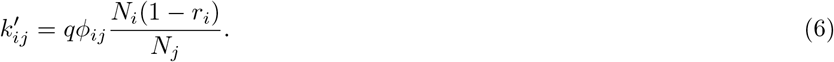

In classical mathematical epidemiology in a well-mixed population, the relationship between the basic reproduction number *R*_0_ and the effective reproduction number *R* is

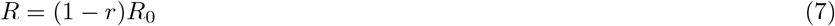

where *r* is the proportion of the population that is immune. We interpret

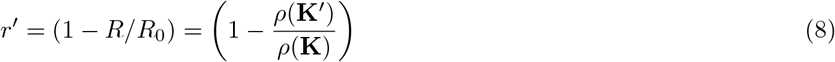

as *contact-adjusted* immunity, that is the equivalent of population immunity once age-specific contact patterns are taken into account. Note that in *q* cancels out, so that calculation of contact-adjusted immunity only requires the contact matrix *φ_ij_*, population sizes *N*_*i*_ and immunity levels *r*_*i*_.

An assumption of homogeneous mixing is equivalent to assuming that *φ_ij_* = *δn_j_*, that is the rate of contact of group *i* being with group *j* depends only on an overall level of contact *δ* and the proportion *n*_*j*_ = *N_j_/N* of the population that are in group *j*, *N* = ∑*N*_*j*_ being the overall population size. This, in turn, means that the infection rate is *β_ij_* = *δp*_inf_*n*_*j*_ and the force of infection (Eq. 1) is independent of age group:

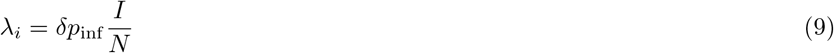

This is equal to the force of infection in a standard SIR model with infection rate *β* if we set *β* = *δp*_inf_, that is the infection rate is equal to the rate of contact times the probability of infection upon contact between a susceptible and infectious individual.

In that case the NGM of Eq. (4) reduces to

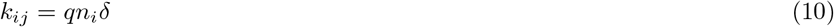

with *q* = *p*_Inf_ *D*_Inf_. This matrix has rank 1 (as all rows are equal), and its only non-zero eigenvalue is given by the trace:

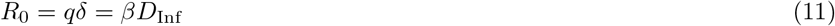

If the proportion immune of those in age group *i* is *r*_*i*_, the elements of **K**′ are

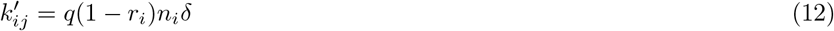

and

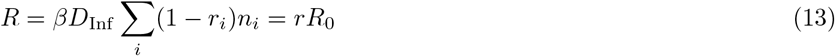

where *r* is the proportion of the population that is immune. We call this factor *r plain* immunity.

### *R*_0_ model: fixed vs. scaled

Elimination is equivalent to a situation where *R <* 1 in Eq. 5. For a given basic reproduction number *R*_0_, this corresponds to contact adjusted immunity *r* being greater than a threshold level *r^∗^* in Eq 8.

The value of the basic reproduction number *R*_0_ would be expected to vary between settings, and this could be reflected in different values across countries [31]. Differences in contact patterns (due to factors such as cultural difference, schooling, population density or demography) would be expected to underlie such differences. It is unclear, though, whether these differences are measurable in diary studies, or whether it is masked by inherent uncertainty in these observations, as well as differences in study design and data collection. We therefore tested two interpretations of the contact matrices estimated by diary studies in order to establish this threshold. Under the first, more conservative interpretation (*fixed R*_0_), the contact matrices were taken to capture differences in contact rates between age groups, but not differences between overall levels of contact between the countries. This is equivalent to setting *R*_0_ to be equal across countries while still allowing difference in the relative contact rates between age groups. In this case, we would expect a single threshold level of contact-adjusted immunity given by *r^∗^* = 1 *−* 1*/R*_0_.

Under the second interpretation (*scaled R*_0_) we assumed *R*_0_ to scale according to the observed contact patterns in each country. In this case, every country would be expected to have a different threshold of contact-adjusted immunity depending on its value of *R*_0_, reflecting the average basic reproduction number within the country.

We calculated a scaling factor *c* for each country such that the basic reproduction number in the country was given by

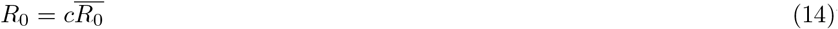

where 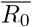 is the mean basic reproduction number across countries. The factor *c* can be calculated as the spectral radius of a given contact matrix divided by the mean of the spectral radii across countries. Instead of working with different values of the basic reproduction number *R*_0_, we rescaled contacted-adjusted immunity in each country as

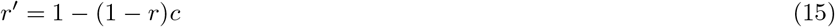

With this formulation, we would again expect a single threshold of scaled contact-adjusted immunity given by 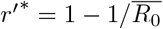.

### Vaccination model: projected vs. ignored

Seroprevalence studies only provide a single, cross-sectional snapshot of immunity in a population. Following such a study, vaccination uptake, natural immunity and ageing combine to change the age-specific immunity levels. We compared a model where vaccination was *ignored* and the measured seroprevalence taken as fixed over the 10-year time period to one where we used an average of *projected* immunity levels, which were updated using information on vaccination uptake in the years following the seroprevalence study. In principle, updating immunity levels with measured vaccination coverage and wild-type measles circulation should improve estimates of population-level immunity. In practice, this relies on accurate measurements of both vaccination coverage and case numbers as well as modelling decisions on assumed vaccine efficacy, maternal immunity and distribution of multiple doses (e.g, randomly vs. preferentially to children that have already received a dose), which could mask any gains made from having up-to-date immunity estimates.

Here, we focused on added immunity due to vaccination and assumed that the added immunity due to wild-type measles circulation was negligible. Serological samples from under-1 year olds were only available from 7 of the 17 countries in the ESEN2 study, and the number of samples from each country too small to produce good estimates (676 samples in total), we combined all these samples to produce an overall estimate of maternal immunity of approximately 40% amongst under-1-year olds. We assumed that immunity in the age group that contained the scheduled age of the first dose of measles was given by a country-specific scaling factor multiplied with the reported coverage in that year. This factor would reflect the proportion of children in that age group immunised at any point in time, as a fraction of the ones immunised by the time of departure from the age group. The factor was estimated by comparing the observed seroprevalence with the level of overage reported in that year. For any subsequent doses, we assumed that the vaccine was preferentially given to those that had received a previous dose or doses of the vaccine, as could be estimated from the reported coverage at the time children in that cohort would have been eligible for the previous dose(s). We assumed a vaccine efficacy per dose of 95%. [32]

### Contact matrices

We established contact matrices from diary studies conducted in a range of different settings using a bootstrap, randomly sampling *P* individuals with replacement from the *P* participants of a contact survey. We then determined a weighted average *d*_*ij*_ of the number of contacts in different age groups *j* made by participants of each age group *i*, giving weekday contacts 5*/*2 times the weight of weekend contacts. We further obtained symmetric matrices, i.e. ones fulfilling *c_ij_n_i_* = *c_ji_n_j_* by rescaling

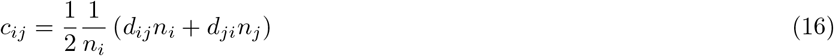

This gave the elements of the contact matrix *φ_ij_* = *c_ij_/T*, scaled by the time period *T* over which contacts were measured (usually 24 hours).

### Data sources

We considered the annual number of measles cases reported by each country to WHO. We used serological studies conducted in 17 countries of the WHO EURO as part of the European Sero-Epidemiology Network 2 (ESEN2) project to determine immunity levels at the times of the studies [10]. Equivocal samples were interpreted as positive as in the original study, but we also tested scenarios where they were removed from the sample or interpreted as negative. We took into account uncertainty by drawing from the individual samples using a bootstrap (*n* = 1000), and using the re-sampled immunity levels with re-sampled contact matrices to estimate contact-adjusted immunity. We ensured visually that the number of bootstrap samples chosen produced stable mean estimates of contact-adjusted immunity levels (see Supplementary Fig. 1). Since contact studies were not available for all countries in ESEN2, contact studies from representative countries were used where necessary (for mediterranean countries: Italy; for Eastern European countries: Poland; for Sweden: Finland; for Ireland: Great Britain).

We used diary studies available on the Zenodo Social Contact Data Repository (https://zenodo.org/communities/social_contact_data), to determine contact matrices for 17 countries and the Hong Kong Special Administrative Region of China [33, 34, 35, 36, 37], a study conducted in Uganda [22] and a further study conducted in five countries of South East Asia.

### Computation

All computations were done with the *R* statistical computing language [38]. Contact matrices were calculated using the *contact matrix* function in the *socialmixr* package [39], and contact-adjusted immunity calculated using the *adjust immunity* function in the *epimixr* package [40].

## Results

### Contact-adjusted immunity levels from serological studies

We first tested the ability of nationwide seroprevalence studies to predict the cases in the decade following, using different definitions of population-level immunity. Overall, the 17 countries that took part in the ESEN2 study in the early 2000s reported 59,494 measles cases to WHO in the 10 years following the study. The number of cases experienced by individual countries varied widely (Fig. 1 and Table 1). Slovakia, where measles was declared eliminated in 1999, only reported a total of 2 cases (both in 2004) in these 10 years. Bulgaria, on the other hand, reported over 20,000 cases, largely as part of a large outbreak in 2009/10.

**Figure 1** Maximum number of cases in a year out of the 10 years following the ESEN2 study, in cases per million inhabitants, on a logarithmic scale. Numbers at the top of the bars are the total number of cases reported in the year with most cases. The dotted vertical line indicates the threshold delineation between countries that did (right) or did not (left) experience large outbreaks when testing the ability of population-level immunity metrics to predict either.

**Table 1.** Measles cases in the 10 years following the ESEN2 serological study, and mean estimated population immunity (contact-adjusted or not, with fixed *R*_0_ and equivocal samples interpreted as positive) based on the study and adjusted for vaccination uptake.

| Country | Cases |  |  |  | Immunity |  |
| --- | --- | --- | --- | --- | --- | --- |
|  | Total (10 years) | Maximum annual | Mean annual (per million) | Maximum annual (per million) | contact-adjusted | plain |
| Slovakia | 2 | 2 | 0.037 | 0.37 | 0.96 | 0.96 |
| Hungary | 12 | 5 | 0.12 | 0.5 | 0.95 | 0.95 |
| Czech Republic | 84 | 30 | 0.82 | 2.9 | 0.98 | 0.98 |
| Latvia | 16 | 7 | 0.72 | 3.1 | 0.71 | 0.82 |
| Sweden | 210 | 62 | 2.4 | 7 | 0.94 | 0.94 |
| Lithuania | 58 | 35 | 1.7 | 10 | 0.91 | 0.93 |
| Slovenia | 26 | 22 | 1.3 | 11 | 0.96 | 0.96 |
| Malta | 14 | 6 | 3.5 | 15 | 0.93 | 0.95 |
| Luxembourg | 10 | 8 | 2.3 | 18 | 0.95 | 0.96 |
| United Kingdom | 6001 | 1445 | 10 | 25 | 0.92 | 0.96 |
| Spain | 3419 | 1842 | 8.6 | 46 | 0.95 | 0.98 |
| Belgium | 1066 | 576 | 10 | 56 | 0.85 | 0.94 |
| Cyprus | 111 | 90 | 11 | 87 | 0.84 | 0.93 |
| Ireland | 1687 | 443 | 40 | 105 | 0.88 | 0.91 |
| Israel | 1792 | 931 | 30 | 155 | 0.94 | 0.95 |
| Romania | 20570 | 7450 | 93 | 337 | 0.92 | 0.96 |
| Bulgaria | 24416 | 22004 | 305 | 2750 | 0.82 | 0.88 |

Comparing the immunity levels with the mean number of annual measles cases in the 10-year period yielded the expected negative correlation with most models (Table 2). Contact-adjusted immunity levels estimated based on the serological profiles were better correlated with the case load than plain immunity levels. Further, interpreting equivocal samples as positive yielded the best correlation, but scaling *R*_0_ according to measured contacts did not improve correlations compared to using a fixed *R*_0_. Projecting national vaccination uptake in the years following the serological surveys onto the observed immunity levels yielded better correlations than just using the snapshots of seroprevalence. For the remaining analyses, we therefore used a fixed *R*_0_, interpreted equivocal samples as positive and corrected immunity levels with vaccination uptake. The resulting immunity levels for the 17 countries in the ESEN2 study are shown in the rightmost two columns of Table 1.

**Table 2.**
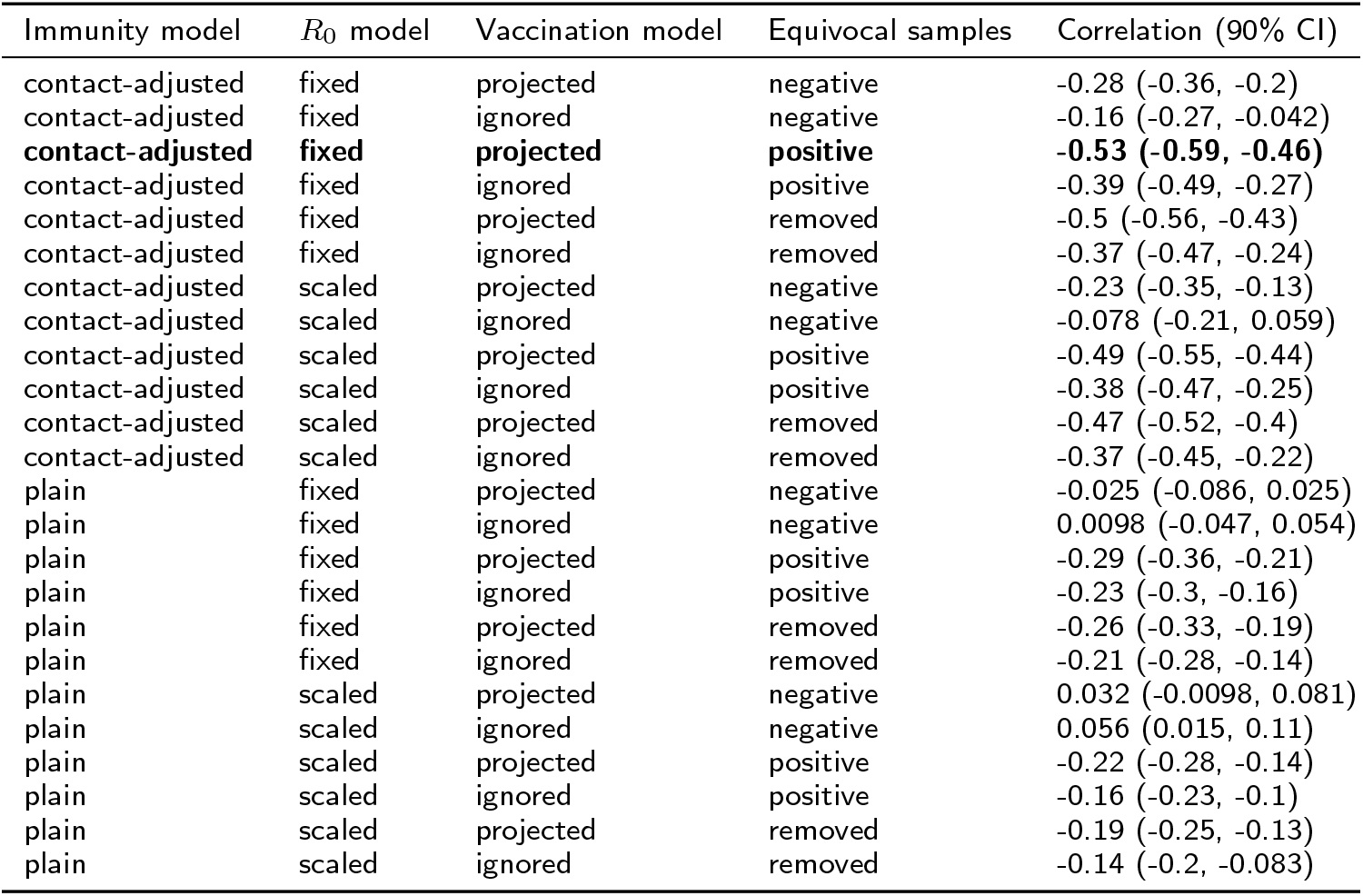
Spearman’s rank correlation between immunity estimated from nationwide serology and (if contact-adjusted) contact studies on the one hand and the mean number of cases in the 10 years following the studies on the other. The model with the greatest absolute correlation is highlighted in bold.

The best model had correlation of −0.53 (Spearman’s rank correlation, 90% credible interval (CI) −0.59–(−0.46)) for contact-adjusted immunity and −0.29 (90% CI −0.36–(−0.21)) for plain immunity. Notable outliers in the correlation between immunity levels and case load were Latvia (contact-adjusted immunity 71%, plain 82%, 16 cases over 10 years) in one direction (low estimated immunity but no outbreaks), and Spain (contact-adjusted immunity 95%, plain 98%, *>* 3000 cases) and Israel (contact-adjusted immunity 94%, plain 95%, *>* 1500 cases) in the other (high estimated immunity but outbreaks).

To test the predictive ability of estimated seroprevalence levels in combination with age-specific mixing we split the countries into those that experienced large outbreaks in the 10 years following the serological studies and those that did not. We set the threshold at an average of 5 per million or, equivalently, a maximum annual cases of 20 per million or (see dashed line in Fig. 1). We then tested different threshold immunity levels (ranging from 80% to 99%, in increments of 1%) and classified countries as being at risk of outbreaks or not based on whether their estimated immunity levels fell below the threshold or not.

The threshold of contact-adjusted immunity yielding best predictions was 93%, in which case about 70% of countries were correctly classified (Fig. 2). With plain immunity this level is at 95%, and the corresponding MCE is greater than with contact-adjusted immunity. More generally, the behaviour of the MCE as a function of threshold level was more erratic when considering plain instead of contact-adjusted immunity. In assessing elimination prospects below, we used the threshold value of 93%.

**Figure 2** Misclassification error (MCE) as a function of the threshold level of for contact-adjusted or plain immunity. Dots give the mean MCE at the tested threshold levels, connected by a line to guide the eye. The grey shades indicate a standard deviation around the mean (uncertainty coming from both the serological sample and from the contact sample).

### Scenarios

We investigated contact-adjusted immunity under previously recommended target immunity levels (85% in under-5 year olds, 90% in 5–9 year olds and 95% in all older age groups) in the settings for which we had access to contact studies (17 countries and Hong Kong, Fig. 3A). We used the identified threshold level of 93% as an indicator of being at risk of outbreaks, implying that the mean *R*_0_ value we found predictive of outbreaks in Europe was a good estimate elsewhere. In this scenario, 5 out of 18 settings had greater than 10% probability of adjusted immunity levels lower than the 93% level found to best identify countries at risk of outbreaks: Taiwan (probability 95%), The Netherlands (90%), Peru (68%), Uganda (63%) and the United Kingdom (40%).

**Figure 3** Contact-adjusted immunity in different theoretical scenarios, with age-specific mixing as measured in diary studies. Each column represents one of the scenarios of age-specific immunity (top), with differences between the settings given by their different mixing patterns. Scenarios from left to right: A) Current target levels. B) 5% higher immunity in under 5 year olds. C) 5% higher immunity in 5–9 year olds. D) 5% lower immunity in 10–14 year olds. E) 5% higher immunity in 5–9 year olds and 5% lower immunity in 15–19 year olds.

With alternative scenarios, the reproduction numbers changed (Fig. 3B–E). Raising immunity in under-5-year olds by 5% to 90% would increase adjusted immunity levels only slightly, with 4 out of the 5 countries (exception: Uganda) at risk under current target immunity levels still at greater than 20% risk. On the other hand, raising immunity in 5-to-9-year olds by 5% to 95% would sharply increase contact-adjusted immunity. In this scenario, all countries would have 5% or less probability of being at risk of outbreaks, with 16 out of 18 at less than 1% risk (exceptions: Hong Kong 5%, Netherlands 3%).

In scenarios where immunity in 5-to-9 year olds was raised but a gap in immunity was introduced in older generations, contact-adjusted immunity dropped below the threshold level of 93% in some settings. A scenario of reduced immunity in 10-to-14-year olds by 5% to 90% while retaining higher immunity in younger age groups resulted in elevated risks of outbreaks in 13 out of 18 countries. A scenario of reduced immunity in 14-to-19 year olds by 5% to 90% while retaining higher immunity in younger age groups resulted in elevated risks of outbreaks in 11 out of 18 countries.

## Discussion

Taking into account age-specific mixing patterns and applying these to immunity levels observed across Europe, we were better able to predict outbreaks than by considering immunity alone. Combined with previous evidence that observed age-specific mixing improve the accuracy of mathematical models, this suggests that there is a case for taking these into account when interpreting the results of sero-logical studies [27, 28, 29].

A threshold of 93% contact-adjusted immunity was found to best predict out-breaks in the subsequent decade, with approximately 70% of countries correctly assessed to either be facing large outbreaks or not. However, in the absence of any more detailed information on setting-specific basic reproduction numbers, such a threshold will only ever be an approximation. On the other hand, setting-specific parameters are difficult to establish, are subject to method-specific biases and can span a wide range of values [41, 31]. In principle, country-specific reproduction numbers should depend on the frequency and types of contact within the population and should therefore be amenable to measurement in contact studies such as the ones used here. Yet, scaling estimated susceptibility levels with the relative number of contacts reported in each study gave no improved results over the simpler version not using such scaling. In other words, while there probably are differences in *R*_0_ between countries, these do not appear to be identifiable in contact studies. At the same time, the contact studies do have value in giving different weights to different age groups when calculating contact-adjusted immunity depending on their contact patterns. We therefore argue that while the achieved 70% of accuracy in predicting outbreaks is far from perfect, aiming to achieve 93% or greater contact-adjusted immunity in a population is a pragmatic choice that can be informed by measurable quantities, that is age-specific immunity levels and mixing patterns.

Current guidelines on target immunity levels are based on estimates derived almost 20 years ago, and were based on assumed mixing patterns matched to pre-vaccination data from England and Wales. We have used transmission models in combination with recently observed age-specific contact patterns from a variety of European and some non-European settings to assess whether these guidelines are sufficient for achieving measles elimination. We investigated a range of settings with different demographic profiles and cultural contexts: from high-income settings characterised by low birth rates and an ageing population (e.g., Germany or the United Kingdom) to having more (Vietnam) or less (Taiwan) recently undergone the demographic transition to low birth rates, or characterised by a high birth rate and young population (Uganda).

With observed mixing patterns, several settings were found to be at risk of out-breaks even if they achieved previously recommended target immunity levels, including ones with very different demographic profiles. Achieving 95% immunity in 5-to-9 year olds, on the other hand, would reduce transmission sufficiently to achieve elimination in all except the most extreme scenarios.

The importance of immunity levels in 5-to-9 year olds presents both a challenge and an opportunity: Levels as high as 95% in this age group can only be maintained through high levels of two-dose immunisation prior to school entry. At the same time, entering this age group coincides with school entry, which involves a level of organisation that provides the opportunity to both check the immunisation status of children and offer additional vaccinations if necessary. The experience of the Pan-American Health Organization in eliminating measles supports these findings. A key component to interrupting measles virus transmission were periodic ‘follow-up’ vaccination campaigns of pre-school children, timed at 4 year intervals to ensure high immunisation by the time of school entry. [42, 43] Studies in the United States, where measles was eliminated in 2000, suggest that different minimum vaccine coverage levels are required to prevent measles virus transmission among different age groups. [44] School-aged populations accounted for the majority of measles cases between 1976 and 1988, and compulsory vaccination as part of school attendance laws played an important role in reducing measles incidence on the path to elimination. [45] Where there were less stringent vaccination requirements at school entry, more case of measles were observed. [46] Analyses of pre-elimination measles outbreaks in the US indicated that transmission occurred among highly vaccinated school-aged populations, suggesting that higher population immunity levels are needed among school-aged children compared to preschool-aged children. [47] It has been proposed that minimum coverage levels as low as 80% at the second birthday of children may be sufficient to prevent transmission among preschool-aged children in the United States if population immunity is at least 93% among over-5 year olds. [48]

While our results stress the role of 5-to-9 year olds, they also highlight the importance of not having gaps in immunity in older age groups. This is particularly important close to elimination as a lower force of infection pushes cases into older age groups. [49] Given the higher rate of complications of measles when experienced at older age, ensuring immunity among adults will be important not only for interrupting transmission, but also to prevent serious episodes of disease. [50]

Our study has several limitations. The delineation of countries into having experienced outbreaks or not is somewhat arbitrary, if in agreement with a milestone towards measles eradication established by the World Health Assembly [51]. Furthermore, the applied threshold of 93% was found to best distinguish between countries that experienced outbreaks and those that did not, but similar performance would have been achieved with thresholds of 92% and, to a slightly lesser extent, 94%. These values correspond roughly to the commonly used range of 12– 18 for the basic reproduction number. Applying these thresholds would have had strong consequences for the assessment of elimination prospects for different immunity profiles, leading to higher (for a threshold of 94%) or lower (for a threshold of 92%) age-specific immunity targets. Depending on the local situation with respect to measles elimination, a country may therefore decide to apply less or more stringent immunity thresholds. Moreover, population immunity represents past levels of vaccine coverage or natural infection which may not be reflective of the future. For example, immunity may be high just after a major outbreak but such outbreaks could occur again if coverage is sub-optimal. In addition, population migration can change immunity levels in a way that is not captured by vaccination coverage figures. An important caveat is therefore that seeing immunity sufficient to interrupt transmission does not guarantee that elimination is maintained if current levels of coverage are insufficient.

Lastly, we assumed that immunity levels and contact patterns alone are sufficient to predict the expected case load. In reality, numerous co-factors such as sub-national heterogeneity or contact patterns that are not captured in age-specific contact matrices (e.g., household and schooling structures) could have influenced this relationship. In fact, the contact-adjusted immunity levels we estimated from serological studies did not always correctly predict where outbreaks could be expected. On the one hand, Latvia did not experience large numbers of cases in spite of low levels of contact-adjusted immunity. It was among the smallest in our group of countries for which we had serological data available and may have been at lower risk of imported cases. Still, they would have been expected to have seen more cases given the results of the serological studies in 2003 and 2004, respectively. Immunity levels were as low as 76% among all age groups and 62% in 5-to-9 year olds in 2003, but only 16 cases of measles were reported in the 10 years 2004–13. Even with the high rates of vaccination coverage (95% coverage of both first and second dose) over these 10 years, outbreaks would have been expected within the age cohorts with large amounts of susceptibility. To our knowledge, there were no supplementary immunisation activities that could explain the absence of outbreaks. It would be of value to determine whether the country is now at high risk of large outbreaks in spite of having previously interrupted transmission, or whether there were issues with the serological tests conducted at the time.

Israel and Spain, on the other hand, experienced large numbers in spite of high levels of contact-adjusted immunity. Three potential causes for this discrepancy suggest themselves: First, in Spain the samples were collected towards in 1996 when there was an ongoing large measles outbreak, and therefore not reflect population-level immunity in the years following. Second, drops in vaccination coverage as well as vaccination campaigns may have changed the risk of outbreaks during the 10 years following the serological studies, although we found nothing in the publicly available national-level vaccination data to suggest any significant changes. Both Spain and Israel consistently reported 94% first-dose MMR coverage in the years following the seroprevalence studies. Third, serology based on residual and population-based samples may not always be representative of relevant immunity levels. In Spain, a disproportionate number of cases occurred in young adults [11], but there was nothing in the serological data to suggest that this might be expected. Moreover, if those lacking immunity are preferentially in contact with each other because they cluster socially or geographically, outbreaks could occur in these groups, population-level serology might not provide a good estimate of realised immunity levels in outbreak settings. In Israel, outbreaks occurred in orthodox religious communities with very low vaccination coverage. [52] More generally, herd immunity thresholds have been shown to increase if non-vaccination is clustered. [53]

These examples highlight that taking into account heterogeneity is crucial. Our method can be applied to lower levels than countries, such as municipalities or counties. Further sub-national serological and epidemiological studies, particularly in low-income countries at high risk of measles outbreaks, could generate key insights on the relationship between immunity levels, heterogeneity of susceptibility and outbreak risk. [54, 55] At the same time, further studies of contact patterns across settings, combined with models of such patterns where no data have been collected, will make it possible to expand our results to other countries and regions. [56]

## Conclusions

We have shown that combining national measles seroprevalence studies with data on contact patterns increases their utility in predicting the expected case load, and in assessing how close a country is to eliminating measles. Comparing past seroprevalence levels to the case load in the following decade enabled us to establish a threshold level of 93% contact-adjusted immunity that appeared sufficient to ensure elimination. Translating this into target age-specific immunity levels that would be necessary to achieve this level of immunity, we found that greater immunity in 5–9 year olds is needed than was previously recommended. While such high levels can be difficult to achieve, school entry provides an opportunity to ensure sufficient vaccination coverage. Combined with observations of contact patterns, further national and sub-national serological studies could serve to highlight key gaps in immunity that need to be filled in order to achieve national and regional measles elimination.

## Declarations

### Acknowledgements

We would like to thank the ESEN2 group for sharing serological data, and the SMILI group for contact data. We further acknowledge fruitful discussions with members of the World Health Organization Strategic Advisory Group of Experts on measles and rubella.

## Availability of data and materials

Contact data are available from the Zenodo Social Contact Data Repository (https://zenodo.org/communities/social_contact_data), except the data for Uganda which is available from the original publication (le Polain de Waroux et al., 2018), and the data for 5 South East Asian countries (Cambodia, Indonesia, Taiwan, Thailand, Vietnam) which are based on the Social Mixing for Influenza Like Illness (SMILI) project. For access to these data, researchers should contact John Edmunds (john@) or Jonathan Read. National-level measles case data were downloaded from the World Health Organization (https://www.who.int/immunization/monitoring_surveillance/data/en/). Seroprevalence data are based on the European Sero-Epidemiology Network 2 (ESEN2) project. Aggregate data have been published previously. For access to the individual-level data, researchers should contact Richard Pebody.

All the results in this paper can be reproduced using code at https://github.com/sbfnk/immunity.thresholds

## Authors’ contributions

SF conducted the analyses and wrote the first draft. All authors contributed to subsequent and final drafts. All authors read and approved the final manuscript.

## Ethics approval and consent to participate

Institutional ethics approval was not sought because this is a retrospective study and the databases are anonymised and free of personally identifiable information.

## Consent for publication

Not applicable.

## Abbreviations

ESEN2: European Sero-Epidemiology Network 2
EURO: European Region
MCE: Misclassification error
WHO: World Health Organization

## Additional Files

Supplementary Figure 1

Mean estimated of contact-adjusted immunity as a function of the number of bootstrap samples. Each line represents one country.

