## Supplementary figures and images for "Combining serological and contact data to derive target immunity levels for achieving and maintaining measles elimination"

### Supplementary Figure 1

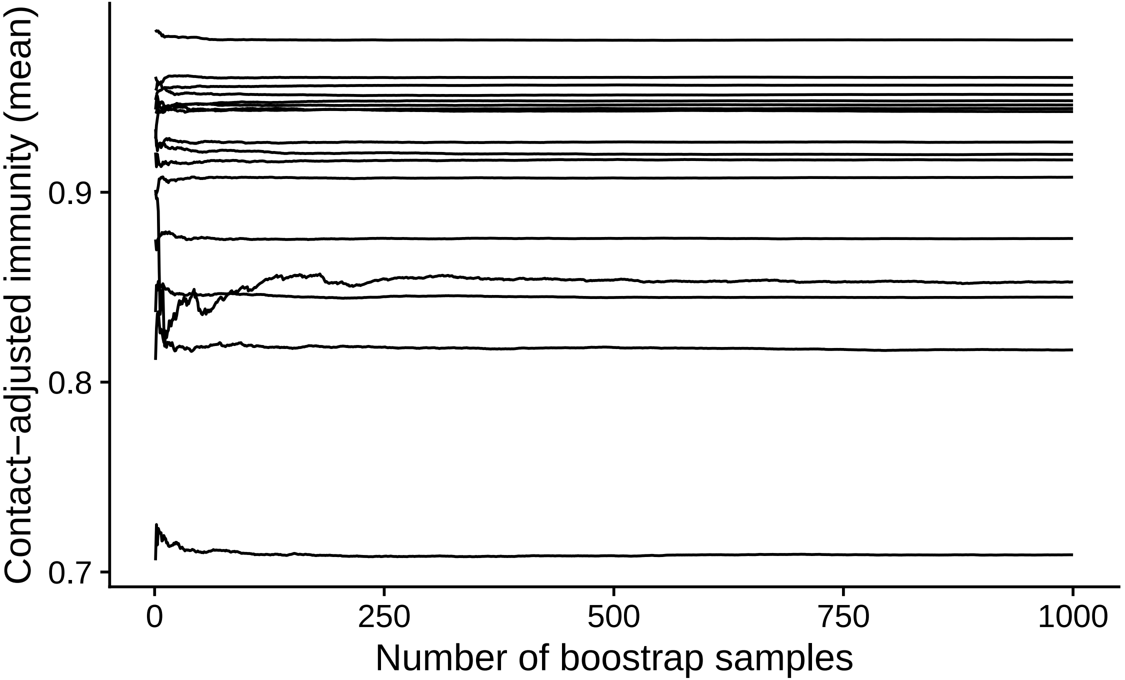
